# Telomere length covaries with age across an elevational gradient in a Mediterranean lizard

**DOI:** 10.1101/732727

**Authors:** Pablo Burraco, Mar Comas, Senda Reguera, Francisco Javier Zamora-Camacho, Gregorio Moreno-Rueda

**Affiliations:** Animal Ecology, Department of Ecology and Genetics, Evolutionary Biology Centre, Uppsala University, Norbyvägen 18D, SE-75236 Uppsala Sweden; Ecology, Evolution and Development Group, Doñana Biological Station (CSIC, Spain); Departamento de Zoología, Facultad de Ciencias, Universidad de Granada (Spain); Estación Biológica de Doñana (EBD-CSIC), Avda. Américo Vespucio 26, Seville E-41092, (Spain); Departamento de Biogeografía y Cambio Global, Museo Nacional de Ciencias Naturales (CSIC, Spain)

**Keywords:** altitude, body condition, ectotherms, growth, reptiles, senescence, telomere shortening

## Abstract

The timing of organisms’ senescence is developmentally programmed but also shaped by the interaction between environmental inputs and life-history traits. In ectotherms, ageing dynamics are still poorly understood despite their particularities concerning thermoregulation, regeneration capacity, or growth trajectory. Here, we investigate the role of life-history traits such as age, sex, body size, body condition, and tail autotomy (i.e self-amputation) in shaping telomere length of six populations of the Algerian sand lizard (*Psammodromus algirus*) distributed across an elevational gradient from 300 to 2500 meters above the sea level. Additionally, we show in a review table the available information on reptiles’ telomere length. We found that telomeres elongated with lizards’ age. We also observed that body size and age class showed a positive relationship, suggesting that cell replication did not shorten lizards’ telomeres by itself. Elevation affected telomere length in a non-linear way, a pattern that mirrored the variation in age structure across elevation. Telomere length was unaffected by tail autotomy, and was sex-independent, but positively correlated with body condition. Our results show that telomeres elongate throughout the first four years of lizards’ lifetime, a process that stress the role of telomerase in maintaining ectothermic telomeres, and, likely, in extending lifespan in organisms with indeterminate growth. Regarding the non-linear impact that elevation had on telomere length of lizards, our results suggest that habitat (mainly temperature) and organisms’ condition might play a key role in regulation ageing rate. Our findings emphasize the relevance of understanding species’ life histories (e.g. age and body condition) and habitat characteristics for fully disentangling the causes and consequences of lifespan trajectory.

## Introduction

The match between phenotypes and the environment not only defines species distribution (Sih, Ferrari, & Harris, 2011; Zamudio, Bell, & Mason, 2016) but also individual lifespan (Monaghan, 2007; Marasco et al., 2017; Ratikainen & Kokko, 2019). The study of the evolutionary underpinnings of ageing has been a long-standing topic both in ecological and medical research, pointing out mitochondrial activity and its underlying damages as the main drivers of individual variation in life expectancy across taxa (Selman, Blount, Nussey, & Speakman, 2012; Ziegler, Wiley, & Velarde, 2015; Vágási et al., 2019). Most studies on ageing in vertebrates have been conducted in endotherms whereas ectotherms have received scarce attention despite their singularities concerning thermoregulation, regeneration capacity, metabolism, or growth trajectory (Olsson, Wapstra, & Friesen, 2018; Monaghan, Eisenberg, Harrington, & Nussey, 2018). Understanding the role of environmental conditions and life-histories on shaping senescence in ectothermic vertebrates would increase the current knowledge about their evolutionary and ecological dynamics.

A reliable way for addressing individual ageing is through estimates of telomere length (Horn et al., 2011; Bateson, 2015; Angelier, Weimerskirch, Barbraud, & Chastel, 2019). Telomeres are non-coding repeated sequences (TTAGGG_n_ in vertebrates) located at the termini of chromosomes, essential for maintaining genomic stability and for protecting cells from chromosome degradation and fusion (O’Sullivan & Karlseder, 2010). Telomeric sequences shorten after each cell replication due to the *end replication problem*, which occurs once the last piece of RNA primer is removed during replication and DNA cannot be extended (Allsopp et al., 1995). Hence, more cell replications may involve shorter length of telomeres. Consequently, telomere length should decrease with age, as often observed in mammals and birds (Haussmann et al., 2003; Heidinger et al., 2012). When telomeres become very short, apoptosis is induced (Aubert & Lansdrop, 2008). However, the expression of telomerase, a reverse transcriptase enzyme that adds repeat sequence to the 3’ end of telomeres, can either partially or fully restore telomere erosion. Telomerase expression is often detected in the germline and in embryonic tissues both in endotherms and ectotherms (Ingles & Deakin, 2016). Particularly in ectothermic vertebrates, telomerase is not only active early in life, but also in adult somatic tissues, like in the fish medaka (*Oryzias latipes*; Klapper, Heidorn, Kühne, Parwaresch, & Krupp, 1998) or in the African water frog (*Xenopus laevis*; Bousman, Schneider, & Shampay, 2003). In this line, telomere elongation has been found throughout larval development of the Atlantic salmon (*Salmo salar*, McLennan et al., 2016) and of the common water frog (*Rana temporaria*; Burraco et al., submitted), and also during the first years of life in some reptiles (e.g. Olsson, Pauliny, Wapstra, & Blomqvist, 2010; Ujvari et al., 2017). In endotherms, telomere elongation after birth is not widespread and takes place under very particular conditions, as during the active season of the edible dormouse (*Glis glis*; Hoelzl, Cornils, Smith, Moodley, & Ruf, 2016), or in some long-lived birds (“elongation hypothesis”, see Haussmann & Mauck, 2007).

Such differences in telomere dynamics between ectothermic and endothermic vertebrates might be linked to organisms’ thermoregulation capacity and growth trajectories (typically, indeterminate growth in ectotherms versus determinate growth in endotherms), and explain lifespan across species (Jones et al., 2014). In reptiles, a paraphyletic group, 11 studies have investigated the variation in telomere length across individuals’ lifetime (Table 1). Four studies found that telomeres shorten with age, whereas in three cases telomere length increased with age in any of the two sexes (Table 1). A quadratic sex-dependent relationship between telomere length and age was observed in two studies, i.e. telomeres increase their length until a certain age, and then shorten (Table 1). Meanwhile, three studies found no effect of age on telomere length in reptiles (Table 1). The high inter-species variation regarding the relation between telomere length across reptiles’ lifetime highlights the need of further research to unravel it.

**Table 1.**
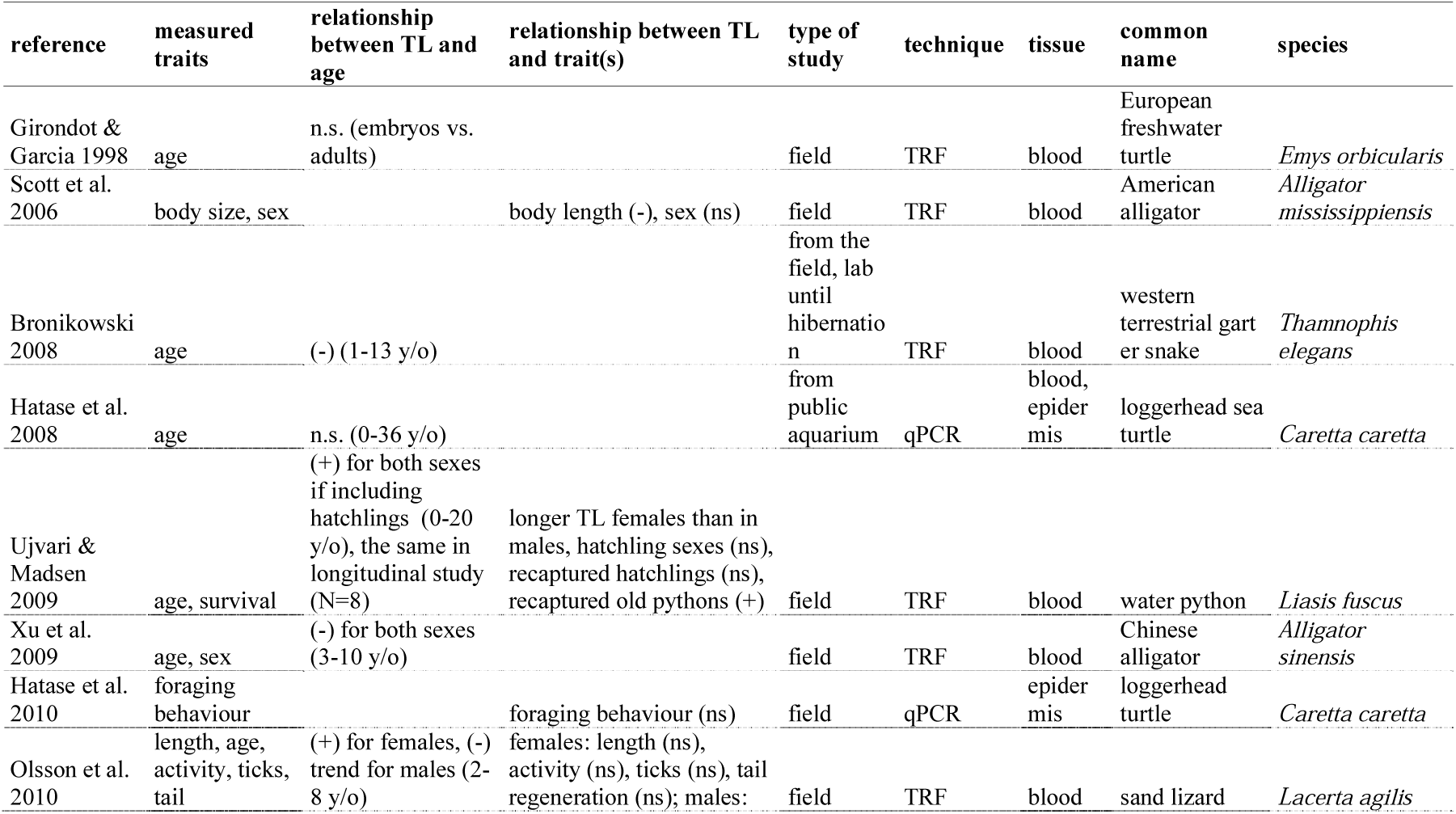
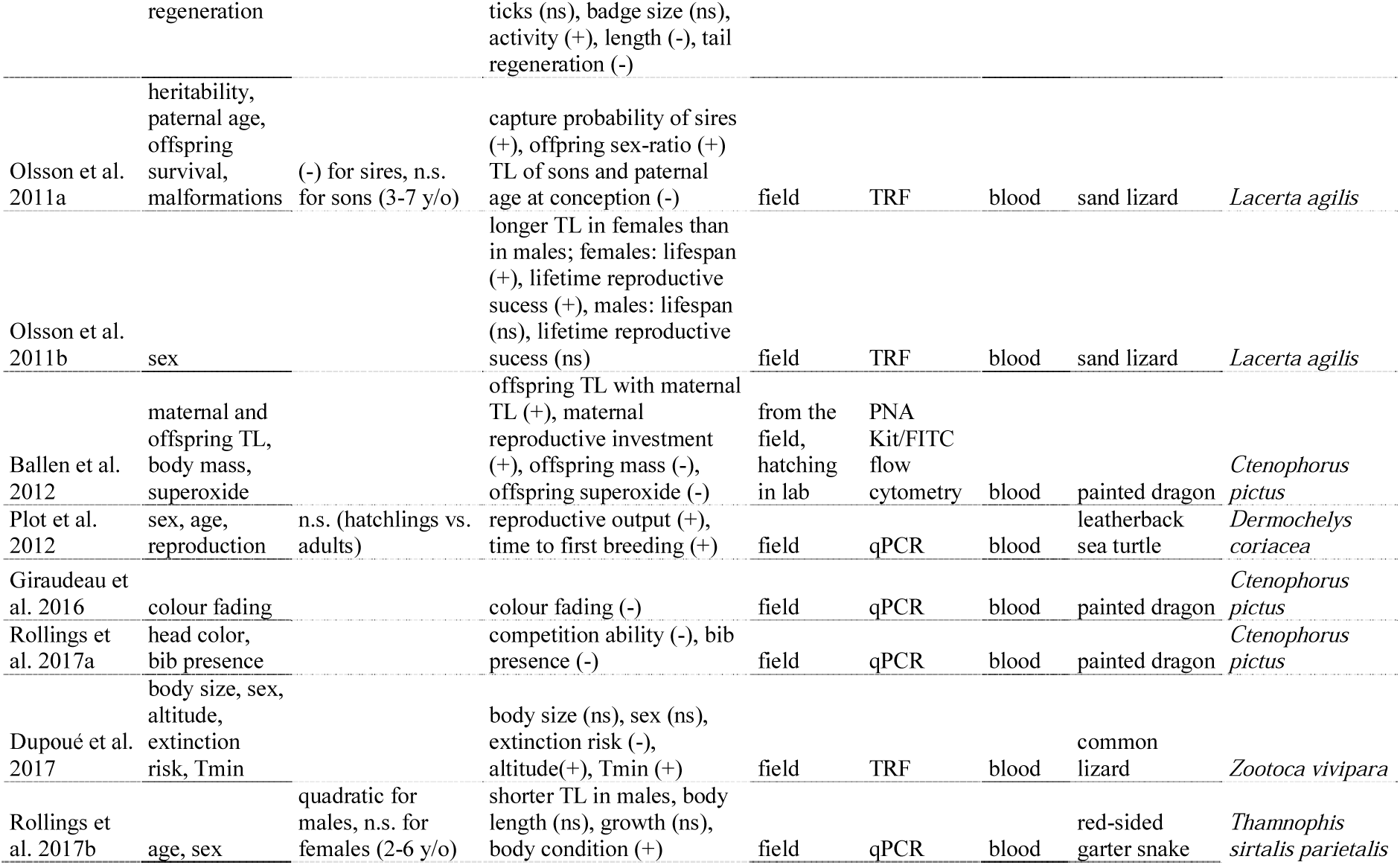
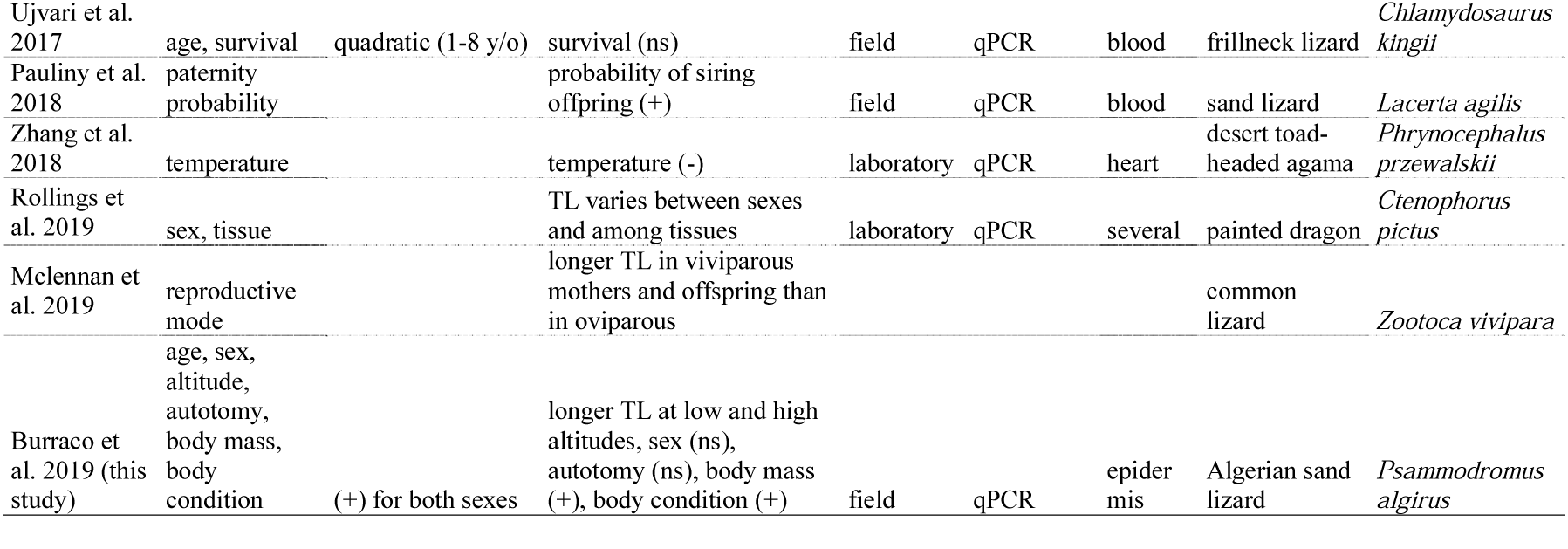
Summary of the studies describing the relationship between telomere length (TL) with age and/or other traits in reptiles. References are included in the Supplementary Information S2.

Telomere length, at a given ontogenetic point, is not only a function of cell replication but also of the organisms’ ability to cope with stress across their life. In vertebrates, harmful conditions often enhance glucocorticoids secretion, which involve metabolic processes that provoke the overproduction of reactive oxygen species (ROS). This overproduction of ROS induces an oxidative state in cells that can damage essential biomolecules like lipids, proteins or DNA (Isaksson, 2015; Luceri et al., 2018), including telomeres (Haussman & Marchetto, 2010; Monaghan, 2014; Angelier, Costantini, Blevin, & Chastel, 2018). As a consequence, telomere length is a reliable indicator of the amount of stress accumulated by organisms across time (Young, 2018). Indeed, positive relationships between telomere length and organisms’ life expectancy (Barret, Burke, Hammers, Komdeur, & Richardson, 2013; Wilbourn et al., 2018), reproductive outcome (Eastwood et al., 2019), or immunocompetence (Alder et al., 2018) are well established. In ectothermic vertebrates, telomere shortening is commonly associated with increased growth rate, bold personality, or predator exposure (reviewed in Olsson et al. 2018). Particularly in reptiles, telomere length positively correlates with social signalling, reproductive output and mode, or lifespan whereas it is unaffected by foraging behaviour or by ectoparasite load (Table 1). One might expect telomeres to shorten as body size increases across lifetime since it implies more cellular replications. However, only a few studies on reptiles have observed a significant effect of body size or growth rate on telomere length (Table 1), unlike in fish (McLennan et al. 2016) or amphibians (Burraco, Díaz-Paniagua, & Gomez-Mestre, 2017a).

Here, we aim to understand the role of life-history traits and environmental conditions on ageing of a lizard species. To this end, we investigated the effect of age, sex, body size, and body condition on telomeres across six populations of the Algerian sand lizard (*Psammodromus algirus*) across a substantial mountain gradient (from 300 to 2500 metres above sea level, m.a.s.l thereafter). We predicted an effect of age class and body size on telomere length, either in a positive, negative, or quadratic way, which may be sex-linked, regarding the available literature on reptile telomeres (Table 1). The elevational gradient studied here allows to determine to which extent environment can influence telomere dynamics in lizard populations. As we ascend in altitude, temperature and activity time decrease while hibernation time increases (Zamora-Camacho, Reguera, Moreno-Rueda, & Pleguezuelos, 2013), which may induce lower telomere attrition (Hoelzl et al. 2016; Kirby, Johnson, Alldredge, & Pauli, 2019). Also, at higher elevations, lizards are exposed to conditions that might reduce telomere shortening such as low risk of overheating (Zamora-Camacho, Reguera, & Moreno-Rueda, 2016) or low ectoparasitism (Álvarez et al., 2018). In addition, oxidative damage –one of the main drivers of telomere attrition (Reichert & Stier, 2017)– decreases at higher elevation (Reguera, Zamora-Camacho, Trenzado, Sanz & Moreno-Rueda, 2014, Reguera et al., 2015). Therefore, we predict that, once corrected for age, populations at higher elevation would have longer telomeres.

## Material and methods

### General procedures

The lizard *P. algirus* is a medium-large lacertid (53-80 mm snout-vent length –SVL-in our study area) that inhabits shrubby habitats in the Mediterranean region from south-western Europe and north-western Africa (Salvador, 2015). In the Sierra Nevada mountain system (SE Spain), we sampled individuals from six populations, which inhabit at 300, 700, 1200, 1700, 2200, and 2500 m.a.s.l. (Fig. 1). In total, we caught 106 individuals (50 males and 56 females): 7 in 2010, 28 in 2011, 65 in 2012 and 6 in 2013. Because lizards were part of a long-term study, we marked individuals by toe clipping, a marking method frequently used in lizards, and that have limited impact on their fitness (Perry, Wallace, Perry, Curzer, & Muhlberger, 2011). We conserved toe samples in ethanol and used them for age class determination using phalanx skeletochronology (more details below). We collected a portion of the terminal region of lizards’ tail (∼ 1 cm) in the field and immersed it in an Eppendorf tube filled with 1.5 mL of absolute ethanol for genetic analyses. Lizards regenerate lost tails, so the small portion we sampled should have had no effects on lizard fitness or welfare. We took special care to disinfect the wounds caused by both toe clipping and tail sampling with chlorohexidine closing the wounds with a tissue adhesive glue (Dermabond®).

**Figure 1.**
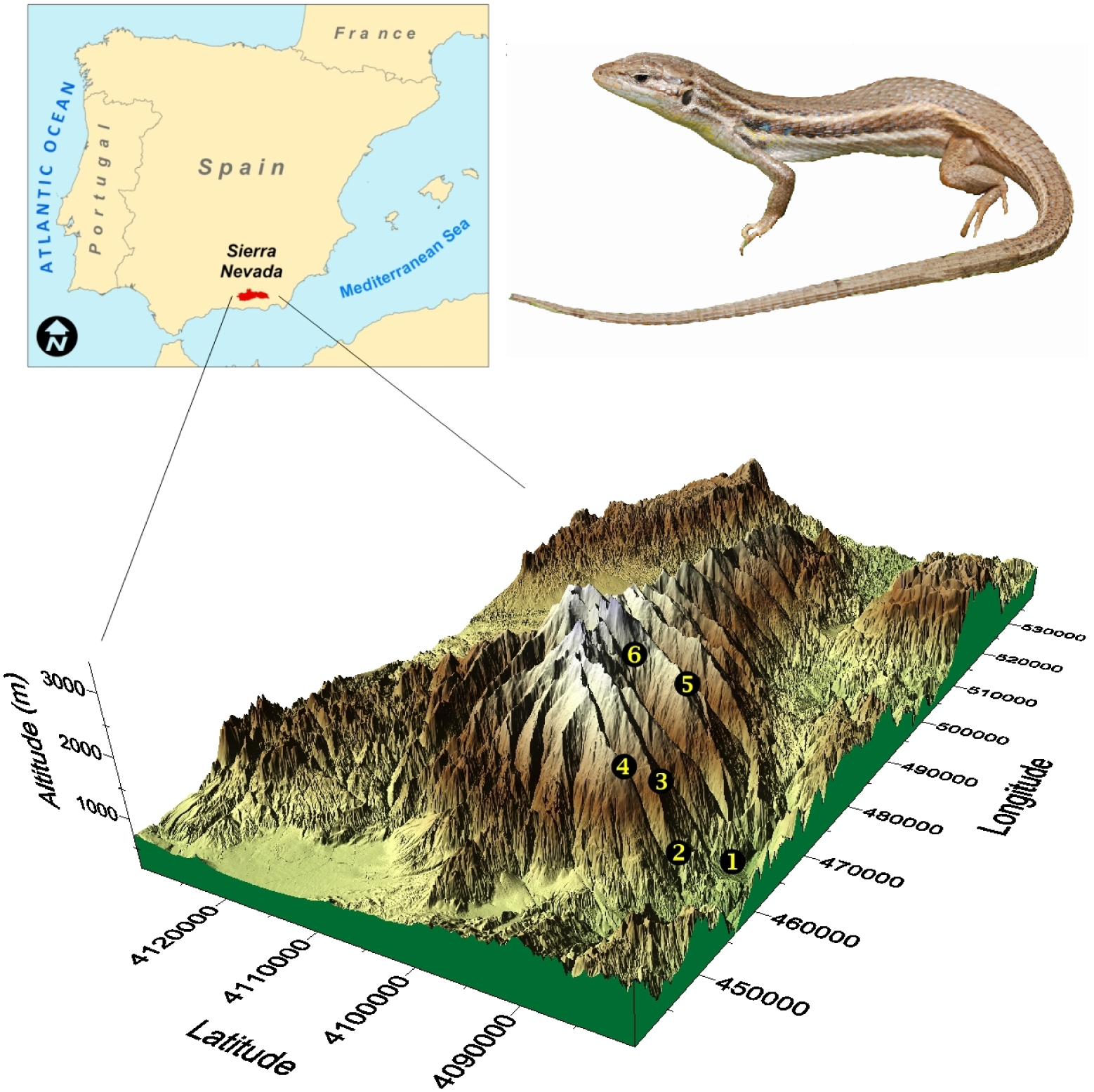
Sampling locations used in this study across an altitudinal mountain gradient (Sierra Nevada mountain system, SE Spain). Numbers from one to six correspond with each location, i.e. 300, 700, 1200, 1700, 2200, and 2500 m.a.s.l. respectively.

We measured lizard body mass with a digital balance (Model Radwag WTB200; to the nearest 0.01 g) and SVL with a metal ruler (to the nearest 1 mm). We estimated the body condition index (BCI) as the residuals of the regressing log mass on log SVL. This widely used index represents the relative energy reserves of an animal (Schulte-Hostedde, Zinner, Milar, & Hickling, 2005). We also recorded whether the tail was intact or regenerated. Males were distinguished from females mainly because they have more femoral pores in their hind limbs (Iraeta, Monasterio, Salvador, & Diaz, 2011) and an orange spot in the corners of their mouths (Carretero, 2002). Gravid females, recognized by palpation of developing eggs inside the trunk, were translated to a lab and placed in individual terrariums (100×20×40 cm) with a heat cable at one end of the cage, switched on three hours a day (11 h-14 h) to allow thermoregulation, indirect access to sun light, and water (in form of aqueous nutritious gel) and food (*Tenebrio molitor* larvae) *ad libitum*. Substrate was bare soil from the study area. We maintained eggs laid in terrariums until hatching. Then, we took a portion of tail of hatchlings for genetic analyses (see below). In order to avoid pseudoreplication, only one neonate per litter (N = 37) was used in the analyses. Females and their neonates were released at the point the female was caught. No lizard died or suffered permanent pain during the study.

### Telomere length measurement

Once in the laboratory, we stored tail samples at −20 °C until assayed. We extracted DNA from epidermis using a high-salt DNA extraction protocol. Since storage conditions, extraction method, or tissue type can affect telomere length measures (Nussey et al., 2014) we used the same conditions for all samples to avoid confounding factors.

We quantified relative telomere length through quantitative polymerase chain reactions (qPCRs), which is one of the most widely used method for estimating telomere length (Nussey et al., 2014). We compared the cycle threshold (C_t_) of telomeric sequences with the C_t_ of a control sequence that is autosomal and non-variable in copy number (Cawthon, 2002, Nussey et al., 2014). As a reference sequence, we amplified GAPDH sequences using 5’-AACCAGCCAAGTACGATGACAT-3′ (GAPDH-F) and 5′-CCATCAGCAGCAGCCTTCA-3′ (GAPDH-R) as forward and reverse primers, respectively. For telomere sequences, we used 5′CGGTTTGTTTGGGTTTGGGTTTGGGTTTGGGTTTGGGTT-3′ (Tel1b) and 5′-GGCTTGCCTTACCCTTACCCTTACCCTTACCCTTACCCT-3′ (Tel2b) as forward and reverse primers, respectively. Conditions of qPCR for GAPDH fragment consisted of 10 min at 95 °C and 40 cycles of 10 sec at 95 °C, 20 secs at 58 °C, and 1 min at 72 °C, and for telomere fragment of 10 min at 95 °C, and 10 secs at 95 °C, 20 secs at 58 °C, and 1 min at 72 °C. We conducted qPCR assays for each gene in separate plates on a LightCycler 480 (Roche) and ran a melting curve from 65 to 95 °C, as a final step in each qPCR to check for specific amplicons. For each sample, we added 20 ng of genomic DNA and used both set of primers at a final concentration of 100nM in a 20 μL master mix containing 10 μL of Brilliant SYBR Green (QPCR Master Mix, Roche). All samples were run in duplicate. Samples with coefficient of variation higher than 5 % were measured again. We calculated qPCR-plates efficiency by including five serial diluted standards in triplicate, obtained from a *golden standard sample* containing a pool of samples from all populations. We calculated the relative telomere length by applying the following formula (Pfaffl, 2001): [(E_telomere_)^ΔCt telomere (control-sample)^]/[(E_GAPDH_)^ΔCt GAPDH (control-sample)^]; where E_telomere_ and E_GAPDH_ are the qPCR efficiency of telomere and GAPDH fragment, respectively; ΔCt telomere (control-sample) and ΔCt GAPDH (control-sample) are the deviation of standard – telomere or GAPDH sequences for each sample, respectively. Efficiencies of qPCR were 1.99 ± 0.02 and 1.93 ± 0.02 for GAPDH and telomere fragments, respectively.

### Estimation of age class with skeletochronology

We estimated individual age class by phalanx skeletochronology (Comas, Reguera, Zamora-Camacho, Salvadó, & Moreno-Rueda, 2016), one of the most accurate techniques to estimate age in many vertebrates, including reptiles (Zhao, Klaassen, Lisovski, & Klaassen, 2019). Vertebrate ectotherms show indeterminate growth, and consequently present a cyclic growth pattern in hard body structures such as bones, corresponding to alternate periods of growth and resting. This pattern is particularly marked in temperate climates, where age can be fairly estimated by counting annual growth rings in the phalanges (Comas et al. 2016). Growth rings are called lines of arrested growth (LAGs). Toes sampled were decalcified in 3% nitric acid for 3 h and 30 min. Cross-sections (10 μm) were prepared using a freezing microtome (CM1850 Leica), stained with Harris hematoxylin for 20 min and dehydrated through an alcohol chain (more details in Comas et al. 2016). Next, cross-sections were fixed with DPX (mounting medium for histology), mounted on slides, and examined for the presence of LAGs using a light microscope (Leitz Dialux20) at magnifications from 50 to 125x. We took several photographs (with a ProgresC3 camera, at the University of Barcelona UB) of various representative cross–sections, discarding those photographs in which cuts were unsuitable for observing LAGs. The number of LAGs detected in the periosteal bone was independently and blindly counted three times by a single observer (MC) on three independent occasions.

### Statistical analysis

We confirmed that the residuals of the models met parametric assumptions (Quinn & Keough 2002). In order not to violate those assumptions, we log-transformed relative telomere length, body mass, and body condition data. We examined the presence of outliers through a Cleveland plot (Zuur, Ieno, & Elphick, 2010), which revealed that an individual had an extremely abnormal low value of relative telomere length, so we decided to omit this data from all the analyses. We performed a linear model to check for differences in telomere length according to the year of capture in order to evaluate possible cohort effects. Since we sampled lizards with intact tail (n = 44) and regenerated tail (n = 58), and tail regeneration could affect the length of telomeres in tail tissue (Anchelin, Murcia, Alcaraz-Perez, Garcia-Navarro, & Cayuela, 2011; Tan et al. 2012; Alibardi, 2016), we tested whether there were differences in relative telomere length between lizards with intact or regenerated tail through linear models. We also performed linear models to examine the effect of sex, age class, and elevation on the relative telomere length. We examined the relationship between relative telomere length and body mass or length, and also between telomere length and age class Pearson correlations. We checked for the independent effect of elevation and age class on relative telomere length by conducting a linear model, in which the effect of sex nor autotomy were not significant and we discarded this variable from the full model. Additionally, we conducted a model selection attending to Akaike information criterion (AIC; Akaike, 1973), and following the recommendations of Burnham & Anderson (2002). The full model contained all measured variables, i.e. elevation, body condition, age class, SVL, sex, tail autotomy, and year of capture. We generated 128 models and calculated the model average of the top 2-AIC models. All statistical analyses were conducted in Statistica software (version 8.0).

## Results

Lizards did not show sexual dimorphism in body mass (F_1, 102_ = 0.11, *P* = 0.74) and sexes did not differ in the distribution of age classes (F_1, 103_ = 1.70, *P* = 0.20). The frequency of male and female lizards did not differ across lizard populations (χ^2^ = 7.35, *P* = 0.69). Relative telomere length did not differ between sexes (F_1, 103_ = 0.30, *P* = 0.59; Figure 2A) and neither with the year of capture (F_3, 101_ = 0.45, *P* = 0.72). The frequency of lizards with autotomized tails did not vary among lizard populations (χ^2^_5_ = 1.36, *P* = 0.93), and tail autotomy did not affect lizard telomere length (F_1, 100_ = 0.00, *P* = 0.99; Figure 2B).

**Figure 2.**
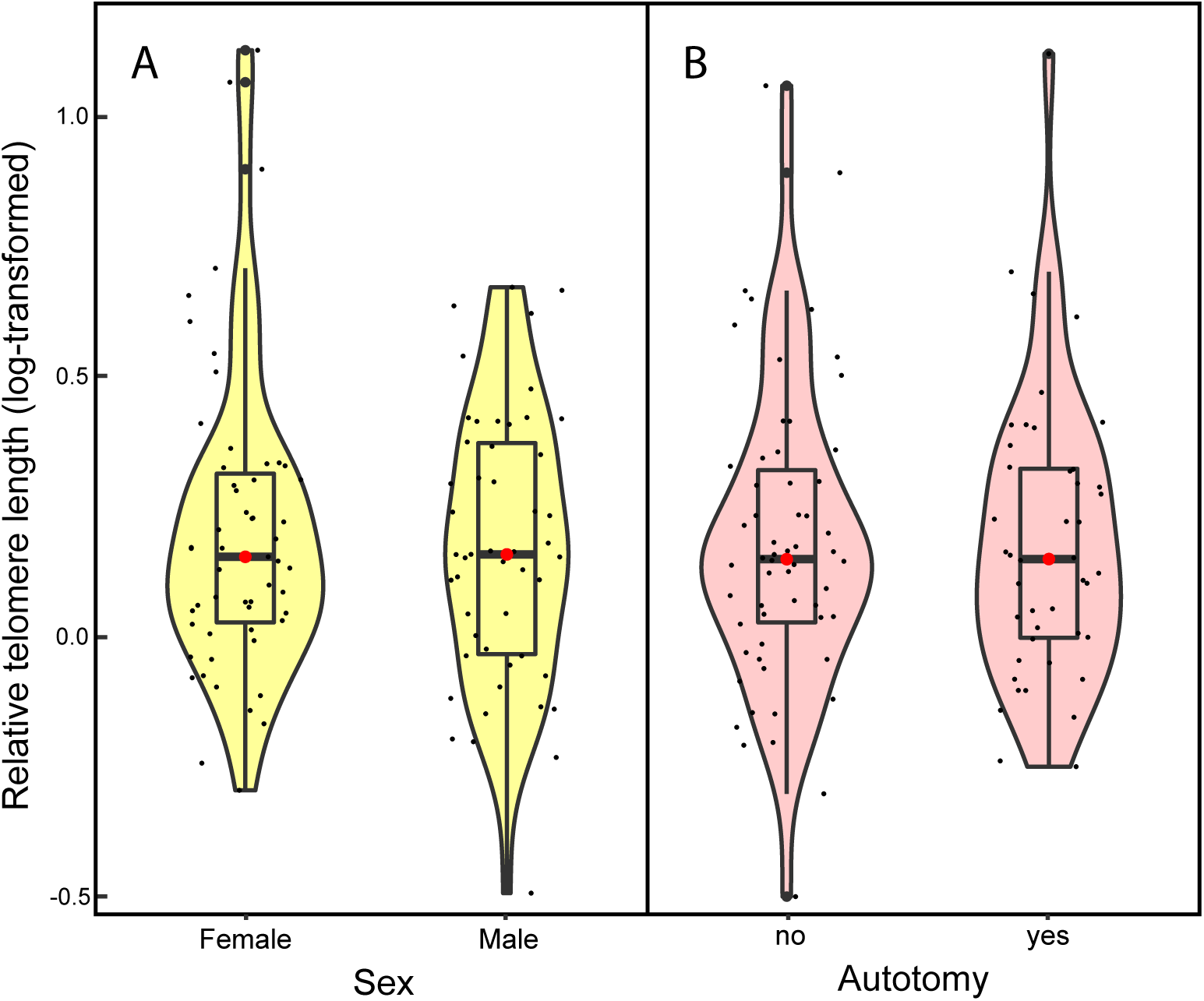
Variation in relative telomere length between sexes (A) or in response to autotomy (B) in individuals of the Algerian sand lizard (*Psammodromus algirus*). The red point shows the mean value at each age and the boxplot the interquartile range. The kernel density plot shows the probability density of data at different values.

Relative telomere length increased with age class, at least until the fourth year (F_5, 136_ = 3.21, *P* = 0.009, Fig. 3). In individuals with five years, telomeres tended to shorten, but we only sampled two lizards with this age class. Spearman correlations between telomere length and age showed similar results (r = 0.24, *P* = 0.003, including neonates, N = 147; r = 0.19, *P* = 0.047, without neonates, N = 105). Age correlated positively with body mass (r = 0.57, *P* < 0.001), which is common in organisms with indeterminate growth like lizards. Likewise, larger individuals had longer telomeres (r = 0.26, *P* = 0.007; Fig. 4), which confirms that telomeres did not shorten by cell replication, but elongated, in those larger and older individuals. Relative telomere length tended to increase with body condition (r = 0.18, *P* = 0.067). This relationship became significant (r = 0.20, *P* = 0.043) when a possible outlier –an individual with very high body condition, indicated by the Cleveland plot– was removed (Fig. 5A). Body condition increased with elevation (F_5, 98_ = 3.03, *P* = 0.014; Fig. 5B).

**Figure 3.**
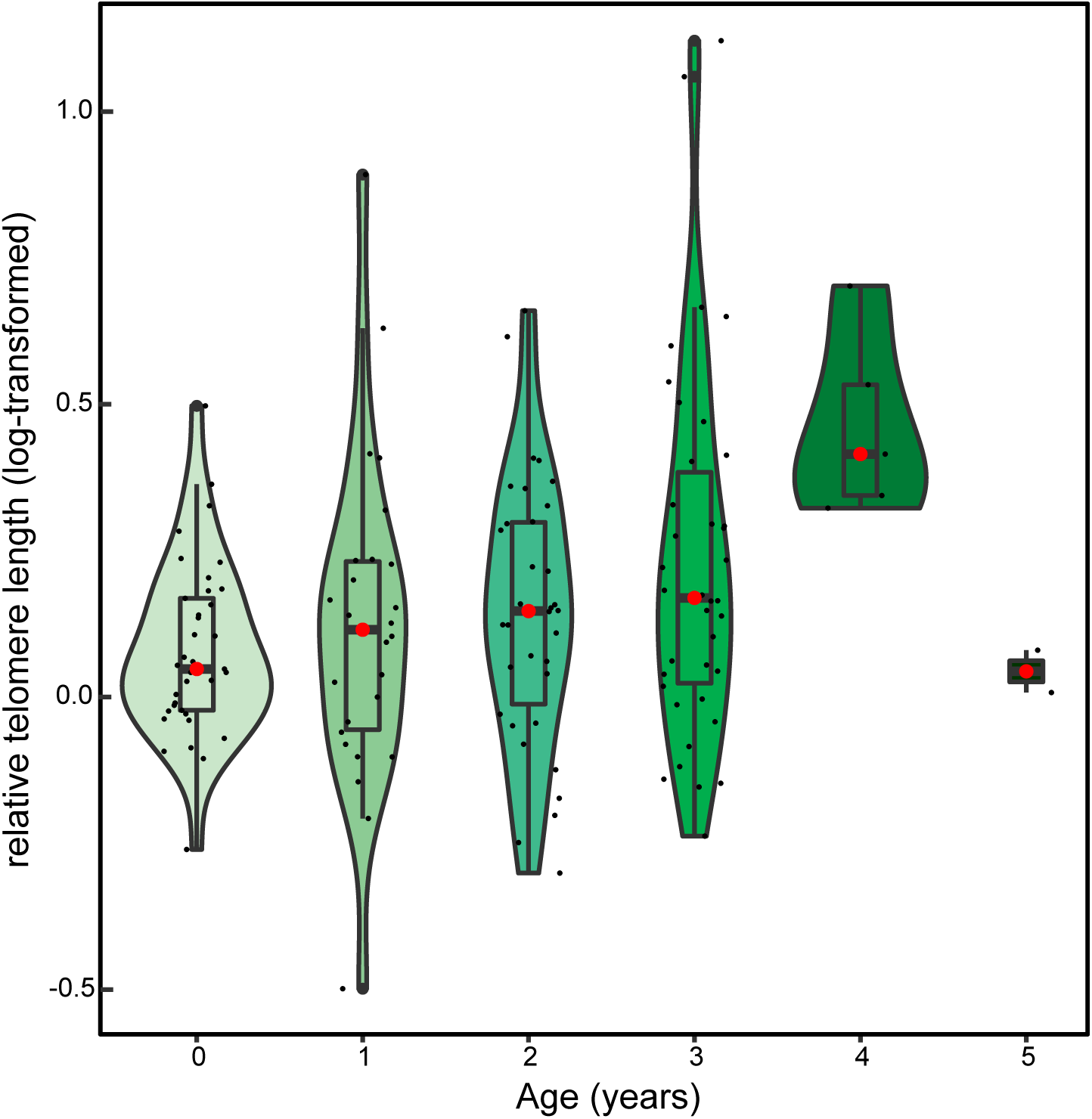
Variation in relative telomere length across lifetime of in individuals of the Algerian sand lizard (*Psammodromus algirus*). The red point shows the mean value at each age and the boxplot the interquartile range. The kernel density plot shows the probability density of data at different values.

**Figure 4.**
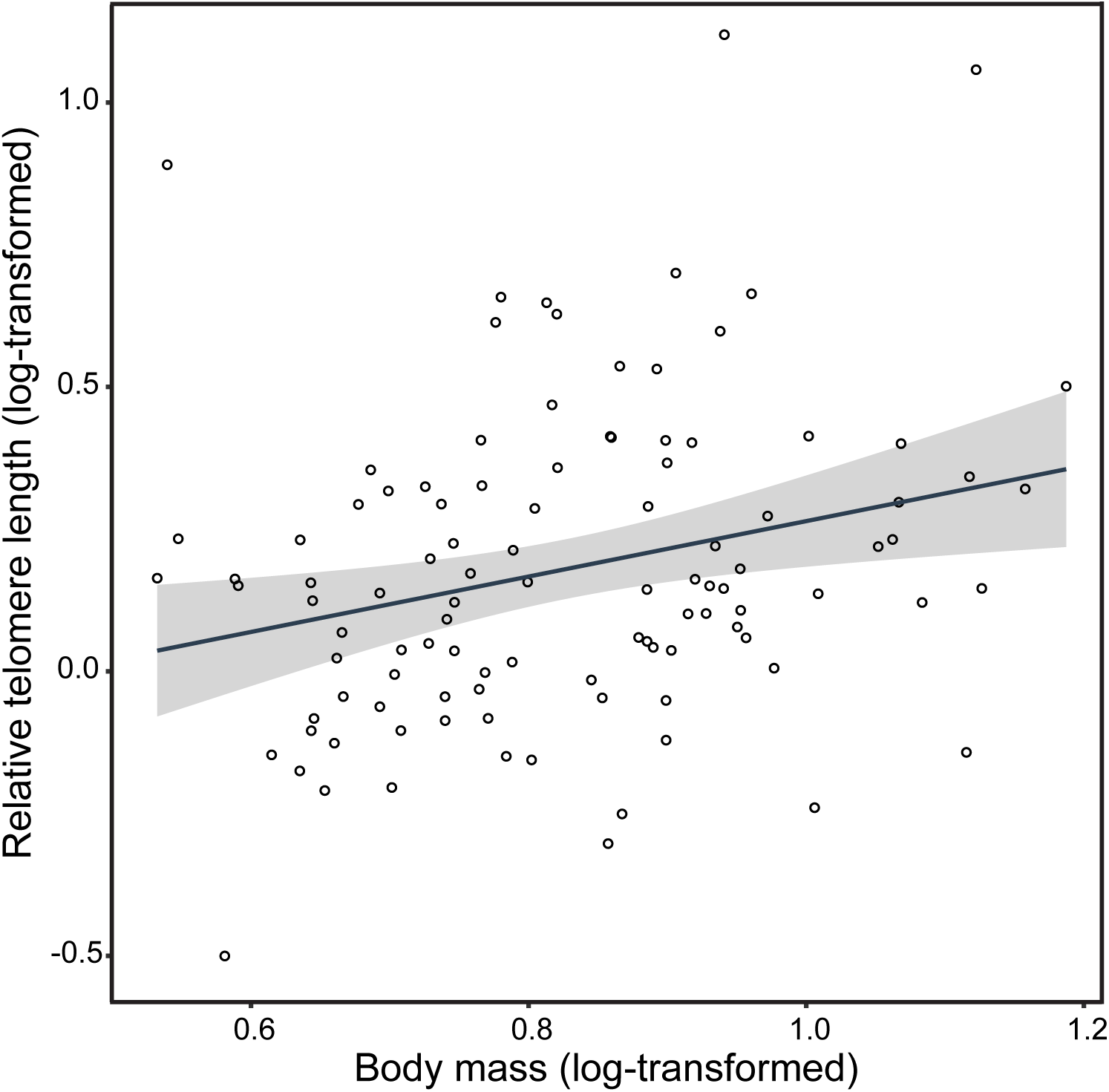
Regression between body mass and relative telomere length (r = 0.26, *P* = 0.007). Regression line shows the correlation between both parameters in all individuals of the Algerian sand lizard (*Psammodromus algirus*) sampled in this study, and indicates that telomeres did not shorten, but elongated with cell replication, as observed in larger (and older) individuals. The grey region indicates the 95% confidence intervals.

**Figure 5.**
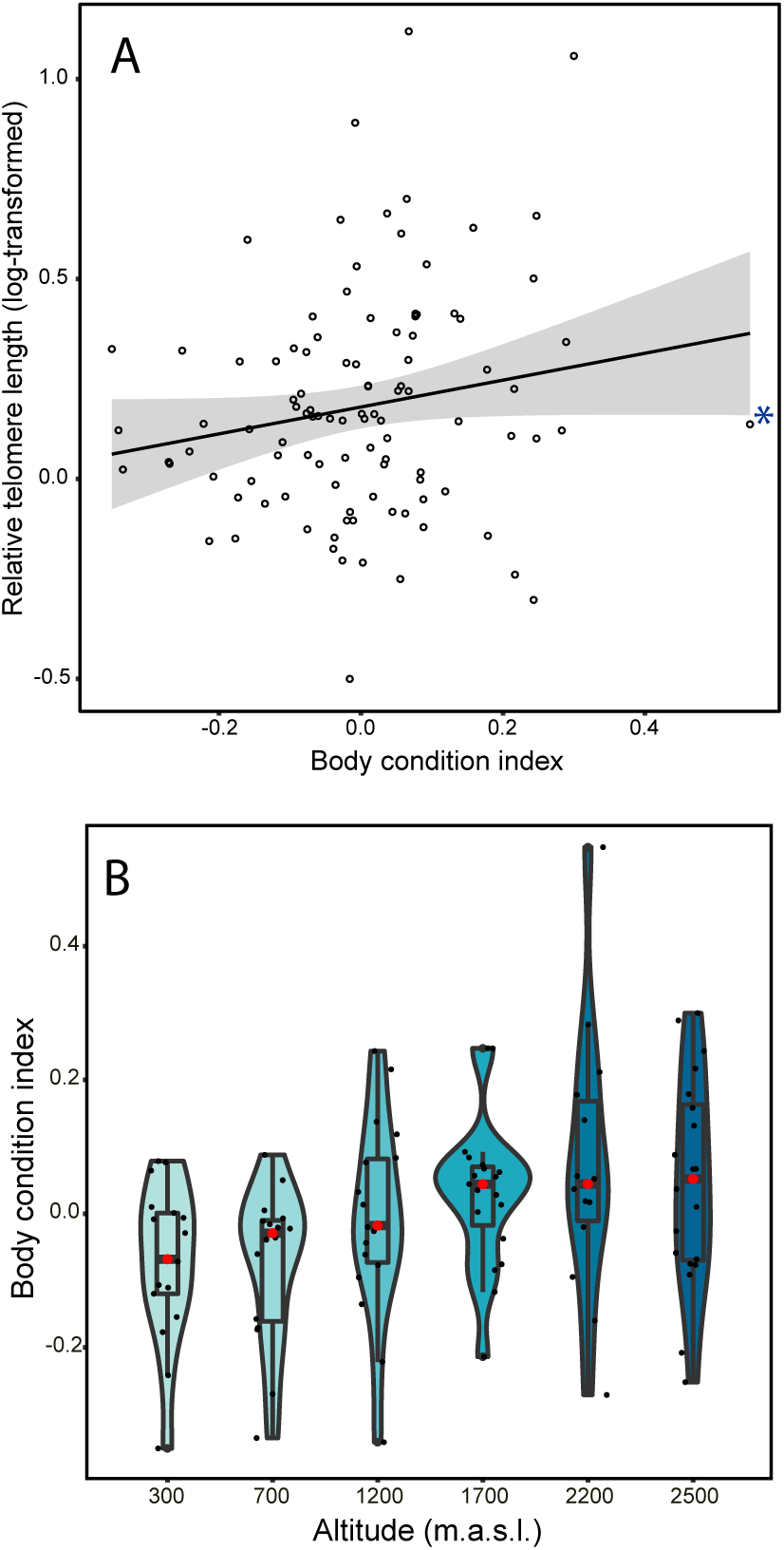
(A) Regression between body condition and relative telomere length (r = 0.20, *P* = 0.043). The asterisk indicates a possible outlier (B) Variation in lizards’ body condition across altitude (F_5, 98_ = 3.03, *P* = 0.014) in individuals of the Algerian sand lizard (*Psammodromus algirus*). The red point shows the mean value at each elevation and the boxplot the interquartile range. The kernel density plot shows the probability density of data at different values.

Lizard telomere length among lizard populations inhabiting across an elevational gradient, but following a non-linear pattern (F_5,136_ = 2.52; *P* = 0.03 for all individuals, and F_5,99_ = 2.07; *P* = 0.070 when excluding neonates; Fig. 6). Individuals at 300 and at 2200 m.a.s.l. had the longest telomeres. However, average age varied with elevation in a similar way (F_5,131_ = 5.44; *P* < 0.001; Figure S1). When we tested the combined effect of age and elevation on telomere length, the effect of age remained significant (F_5,131_ = 2.32; *P* = 0.047), but the effect of elevation was no longer significant (F_5,131_ = 1.67; *P* = 0.15). Model selection showed that body length (SVL) had the highest explanatory power to understand variation in telomere length in our study system, which is presumably explainable by the positive relationship between telomere length and age.

**Figure 6.**
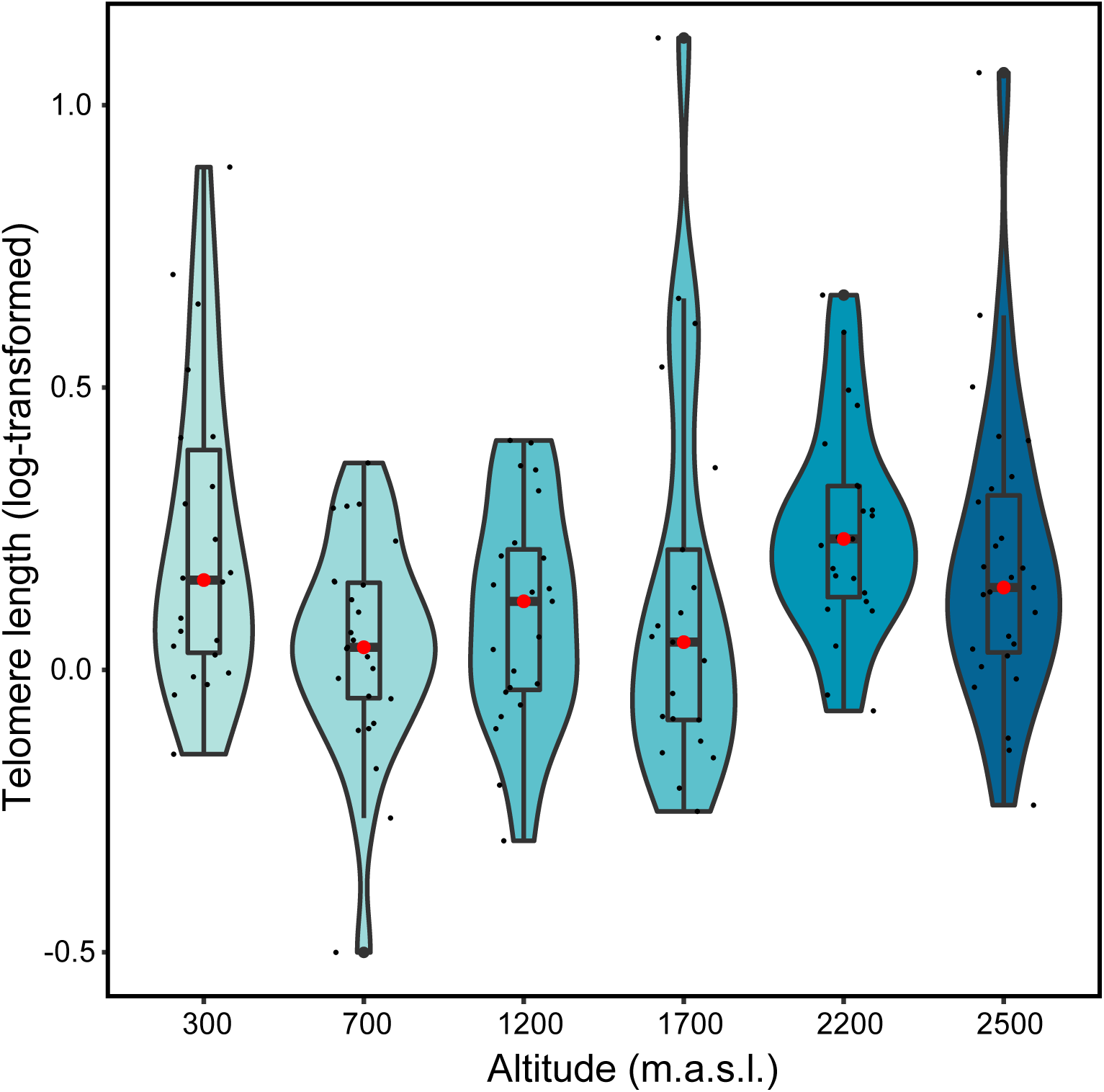
Variation in relative telomere length across altitude in individuals of the Algerian sand lizard (*Psammodromus algirus*). The red point shows the mean value at each age and the boxplot the interquartile range. The kernel density plot shows the probability density of data at different values.

Neonate telomere length, an indicator of the baseline telomere length at birth, varied among populations (F_5,31_ = 2.91; *P* = 0.03), but with no clear pattern; lizard neonates showed the longest telomeres at 2200 m.a.s.l., but the shortest at 1700 m.a.s.l. (Fig. S2).

## Discussion

Life-history trade-offs and environmental conditions can shape ageing across taxa (Wilbourn et al., 2018; Eastwood et al., 2019). Here, we show that, in the Algerian sand lizard, telomeres elongated with age until their fourth year. Additionally, larger lizards had longer telomeres. Intriguingly, although lizard populations across this substantial elevational gradient differed in their telomere length, differences were not linear and the variation in telomere length mirrored the variation in the distribution of age classes across the elevational gradient. On the other hand, telomere elongation was sex-independent, unlike in adults of other sand lizard species (*Lacerta agilis*, Olsson et al., 2011). Sex differences in telomere length may result from sex differences in growth rate, body size, and/or age (Olsson et al., 2018). However, in our study system, lizards did not show sexual dimorphism in size or age structure. Autotomy did not affect telomere length despite the fact that differences in the regulation of telomere length may be driven by evolutionary pressures such as predation (Olsson et al., 2010), and also by enhanced cell replication during tissue regeneration. Moreover, no cohort effect was detected, as telomere length did not differ with year of sampling, which validates our cross-sectional study.

Telomere elongation observed in lizards across their first four years of life agrees with previous studies in snakes and lizards (Ujvari & Madsen, 2009 and Ujvari et al., 2017, respectively). At the fifth year, telomeres tended to shorten, although this result should be interpreted carefully because we only collected two five-year-old individuals. Telomere length showed a positive relationship both with age and body size, suggesting that cell replication does not shorten telomeres by itself. This finding adds to previous studies showing that ectotherms, unlike endotherms, often show telomere elongation along their lifetime (Olsson et al., 2018). Such contrasting pattern of telomere dynamics in ectotherms may be related to a higher telomerase expression after birth in somatic cells in ectotherms than in endotherms (Gomes, Shay, & Wright, 2010). Hence, telomerase may be relevant for buffering downstream effects of ROS in organisms with indeterminate growth such as lizards (Jones et al. 2014). However, telomerase expression may not be enough to protect from telomere shortening in ectothermic vertebrates. For instance, telomerase is expressed in tissues of adult medaka fish (Klapper et al. 1998) but telomeres shorten with age (Hatakeyama et al., 2008). Furthermore, the maintenance of telomerase expression in species with indeterminate growth can imply a trade-off suggested by a higher cancer occurrence in ectotherms (Gomes et al., 2010; Olsson et al. 2018), however, the knowledge about cancer in wildlife is still meagre.

Mountains cover *circa* a quarter of the Earth’s surface (Körner, 2007). Elevational gradients are characterised by deep changes both in biotic and physical conditions, such as competitor and/or predators’ abundance, temperature, or ultraviolet radiation (Barry, 2008). Previous research demonstrated a broad number of physiological adaptations to divergent habitats across altitude (Bozinovic, Calosi, & Spicer, 2011; Keller, Alexander, Holderegger, & Edwards, 2013; Boyle, Sandercock, & Martin, 2016). Such adjustments often imply elevational variation in energy expenditure devoted to reproduction and somatic maintenance, then affecting telomere dynamics (Stier et al., 2016). Also, variation in temperature can involve deep physiological shifts in ectotherms across elevations since their body temperature greatly depends on environmental heat (Angilleta, 2009; Gunderson & Stillman, 2015). In our study, we expected to find shorter telomeres in lizard populations at higher elevation, as we know that higher-altitude lizards undergo reduced activity time and oxidative damage (Zamora-Camacho et al., 2013; Reguera et al., 2014, 2015). However, we found a non-linear variation in telomere length with elevation. The most plausible explanation is that telomere length across elevation mirrored the altitudinal distribution of lizard age. Contrary to our results, Dupoué et al., (2017) found that populations of the common lizard (*Zootoca vivipara*) inhabiting at low elevations have shorter telomeres and higher extinction risk. In our study system, lowland populations also suffer poor habitat quality, such as low thermal quality (risk of overheating, Zamora-Camacho, Reguera, & Moreno-Rueda, 2016), high ectoparasitism (Álvarez et al., 2018), low food availability (Moreno-Rueda et al., 2018), high oxidative damage (Reguera et al., 2014, 2015), and even high risk of wildfire (Moreno-Rueda, Melero, Reguera, Zamora-Camacho, & Comas, 2019). Additionally, at low elevations, lizards increase their activity time while hibernation time decreases (Zamora-Camacho, Reguera, Moreno-Rueda, & Pleguezuelos, 2013). In spite of all this, lizard populations at lowland did not have shorter telomeres than populations at high elevations.

Lizard body condition, temperature, and telomerase expression might have shaped telomere length of lizards inhabiting at different elevations. In this study, body condition of lizards was higher in populations at higher elevation, and correlated positively with telomere length. Telomere length is positively correlated to body condition in other reptiles (*Thamnophis sirtalis*; Rollings et al., 2017). In addition, it is likely a temperature-mediated regulation of telomerase expression, thus at low elevation telomerase might show a higher expression, then compensating for telomere erosion (Olsson et al. 2018). At the highest elevations (mainly at 2200 m.a.s.l.), the reduction in metabolic rate due to cold conditions may have favoured a reduction in the rate of telomere erosion due to a reduced production of ROS, involving an adaptive downregulation of telomerase. Indeed, increases in lifespan are often orchestrated by reductions in metabolic rate (Speakman, 2005), as for example suggested by the straightforward influence of latitude on lifespan of *Rana temporaria* frogs across the Swedish latitudinal gradient (Hjernquist et al., 2012). Furthermore, the variation in the pace-of-life as a consequence of facing particular environmental conditions is also known to alter telomeres, then resulting in complex or unexpected patterns (Giraudeau, Algelier, & Sepp, 2019). For example, shorter telomeres are associated with higher survival in migratory Atlantic salmon (McLennan et al. 2017), thus benefits of experiencing intense telomere erosion can be higher than costs of responding poorly to certain scenarios, such as during migration. Likewise, amphibian larvae surviving predators, which have larger bodies and larger fat reserves, experience telomere shortening as a consequence of growing faster due to relaxed intraspecific competition (Burraco et al. 2017a). In our system, other factors like diseases or intraspecific interactions might have also modulated ageing in lizards. A cross-fostering approach would help to fully clarify the evolutionary impact of both environment and life-history traits on telomeres of this lizard metapopulation.

## Conclusions

Our results show that telomeres elongate throughout the first four years of lizards’ lifetime, a process that stress the role of telomerase in maintaining ectothermic telomeres, and, likely, in extending lifespan in organisms with indeterminate growth. Habitat features and repair mechanisms at different habitats may be relevant for understanding telomere dynamics in ectothermic vertebrates. This study also shows that telomere length can follow a complex trajectory across habitats occupied by an ectothermic vertebrate, as across a substantial altitudinal gradient. Our results emphasize the relevance of understanding species’ life histories (e.g. age and body condition) and habitat characteristics for disentangling the causes and consequences of lifespan trajectory.

## Authors’ contribution

GMR, MC and PB conceived the idea; SR and FJZC performed the sampling; MC and SR carried out the histological labwork and data analysis of skeletochronology; PB carried out the telomere length analysis; GMR and PB performed the statistical analyses; PB wrote the manuscript with inputs from GMR, MC, SR and FJZC.

## Acknowledgements

PB was supported by fellowship F.P.U.-AP2010-5373, and by the Carl Tryggers Foundation project CT 16:344. FJZC (F.P.U.-AP2009-3505) and SR (F.P.U.-AP2009-1325) also were supported by respective fellowships. MC was supported by a Severo Ochoa contract (SVP-2014-068620). This study was economically supported by the Ministerio de Ciencia e Innovación (project CGL2009-13185). The study complies with the current laws of Spain, and were performed in accordance with the Junta de Andalucía and Sierra Nevada National Park research permits (references GMN/GyB/JMIF and ENSN/JSG/JEGT/MCF). We are grateful to Concepción Hernández (Centre of Scientific Instrumentation of the University of Granada) for her help with the freezing microtome, Humbert Salvadó (Universitat of Barcelona) for let us to use his microscopy for this study, and Francisco Miranda (Ecophysiology Laboratory of Doñana Biological Station) for assistance with the telomere assays. We also thank the personnel from the Espacio Natural de Sierra Nevada for their constant support.

## Data accessibility

Data will be accessible at FigShare upon manuscript acceptance.

