## Supplementary figures and images for "Telomere length covaries with age across an elevational gradient in a Mediterranean lizard"

### Supplemental Figure S1

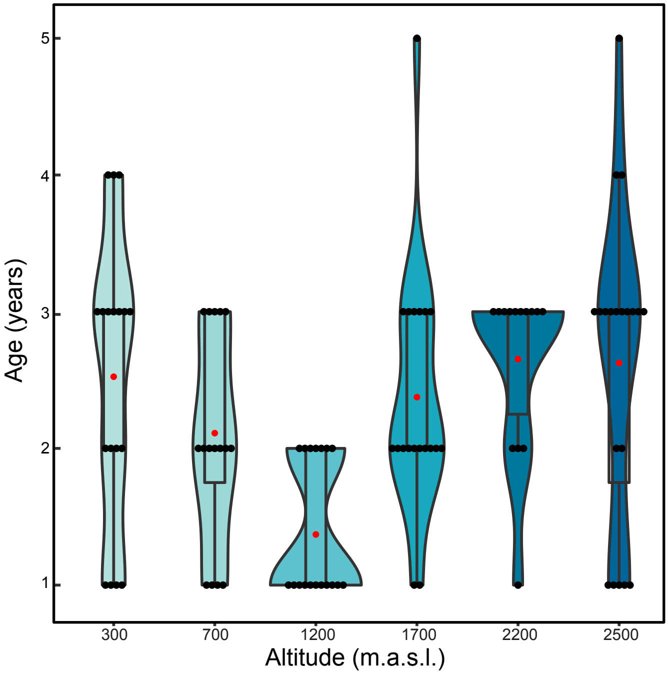

### Supplemental Figure S2

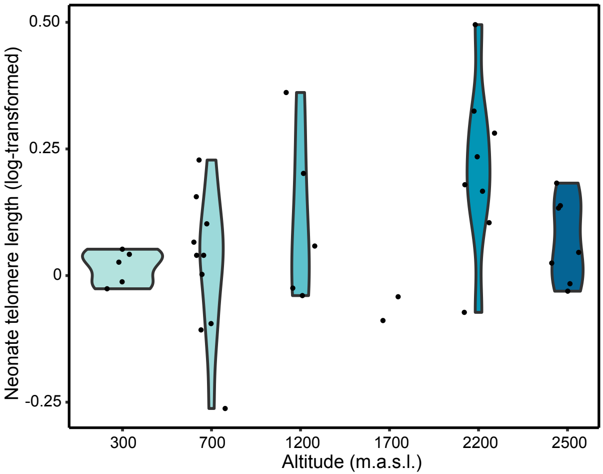
