## Supplemental information for "Telomere length covaries with age across an elevational gradient in a Mediterranean lizard"

**Supplementary Figure S1.** Variation in age across altitude of Algerian sand lizard (*Psammodromus algirus*). The red point shows the mean value at each age and the boxplot the interquartile range. The kernel density plot shows the probability density of data at different values.

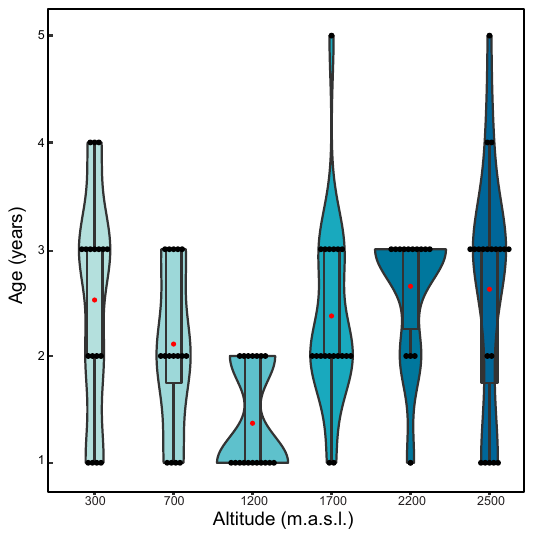

**Supplementary Figure S2.** Variation in relative telomere length across altitude of neonates of the Algerian sand lizard (*Psammodromus algirus*). The red line shows the mean value at each age, whereas the kernel density plot shows the probability density of data at different values.

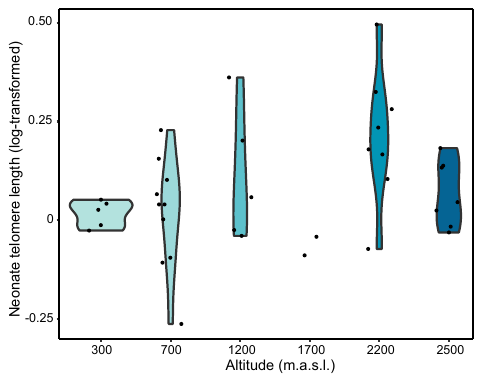

**References Table 1**

Ballen, C., Healey, M., Wilson, M., Tobler, M., & Olsson, M. (2012). Predictors of telomere content in dragon lizards. *Naturwissenschaften*, *99*(8), 661-664.

Bronikowski, A. M. (2008). The evolution of aging phenotypes in snakes: a review and synthesis with new data. *Age*, *30*(2-3), 169-176.

Dupoué, A., Rutschmann, A., Le Galliard, J. F., Clobert, J., Angelier, F., Marciau, C., ... & Meylan, S. (2017). Shorter telomeres precede population extinction in wild lizards. *Scientific Reports*, *7*(1), 16976.

Giraudeau, M., Friesen, C. R., Sudyka, J., Rollings, N., Whittington, C. M., Wilson, M. R., & Olsson, M. (2016). Ageing and the cost of maintaining coloration in the Australian painted dragon. *Biology Letters*, *12*(7), 20160077.

Girondot, M., & Garcia, J. (1999). Senescence and longevity in turtles: what telomeres tell us. In *9th Extraordinary Meeting of the Societas Europaea Herpetologica. Université de Savoie, Le Bourget du Lac, France* (pp. 133-137).

Hatase, H., Sudo, R., Watanabe, K. K., Kasugai, T., Saito, T., Okamoto, H., ... & Tsukamoto, K. (2008). Shorter telomere length with age in the loggerhead turtle: a new hope for live sea turtle age estimation. *Genes & genetic systems*, *83*(5), 423-426.

Hatase, H., Omuta, K., & Tsukamoto, K. (2010). Oceanic residents, neritic migrants: a possible mechanism underlying foraging dichotomy in adult female loggerhead turtles (*Caretta caretta*). *Marine Biology*, *157*(6), 1337-1342.

Mclennan, D., Recknagel, H., Elmer, K. R., & Monaghan, P. (2019). Distinct telomere differences within a reproductively bimodal common lizard population. *Functional Ecology*. *In press.*

Olsson, M., Pauliny, A., Wapstra, E., & Blomqvist, D. (2010). Proximate determinants of telomere length in sand lizards (*Lacerta agilis*). *Biology Letters*, *6*(5), 651-653.

Olsson, M., Pauliny, A., Wapstra, E., Uller, T., Schwartz, T., & Blomqvist, D. (2011a). Sex differences in sand lizard telomere inheritance: paternal epigenetic effects increases telomere heritability and offspring survival. *PLoS One*, *6*(4), e17473.

Olsson, M., Pauliny, A., Wapstra, E., Uller, T., Schwartz, T., Miller, E., & Blomqvist, D. (2011b). Sexual differences in telomere selection in the wild. *Molecular Ecology*, *20*(10), 2085-2099.

Pauliny, A., Miller, E., Rollings, N., Wapstra, E., Blomqvist, D., Friesen, C. R., & Olsson, M. (2018). Effects of male telomeres on probability of paternity in sand lizards. *Biology Letters*, *14*(8), 20180033.

Plot, V., Criscuolo, F., Zahn, S., & Georges, J. Y. (2012). Telomeres, age and reproduction in a long-lived reptile. *PloS One*, *7*(7), e40855.

Rollings, N., Friesen, C. R., Sudyka, J., Whittington, C., Giraudeau, M., Wilson, M., & Olsson, M. (2017a). Telomere dynamics in a lizard with morph‐specific reproductive investment and self‐maintenance. *Ecology and Evolution*, *7*(14), 5163-5169.

Rollings, N., Uhrig, E. J., Krohmer, R. W., Waye, H. L., Mason, R. T., Olsson, M., ... & Friesen, C. R. (2017b). Age-related sex differences in body condition and telomere dynamics of red-sided garter snakes. *Proceedings of the Royal Society B: Biological Sciences*, *284*(1852), 20162146.

Scott, N. M., Haussmann, M. F., Elsey, R. M., Trosclair, P. L., & Vleck, C. M. (2006). Telomere length shortens with body length in *Alligator mississippiensis*. *Southeastern Naturalist*, *5*(4), 685-693.

Ujvari, B., & Madsen, T. (2009). Short telomeres in hatchling snakes: erythrocyte telomere dynamics and longevity in tropical pythons. *PloS One*, *4*(10), e7493.

Ujvari, B., Biro, P. A., Charters, J. E., Brown, G., Heasman, K., Beckmann, C., & Madsen, T. (2017). Curvilinear telomere length dynamics in a squamate reptile. *Functional Ecology*, *31*(3), 753-759.

Xu, M., Wu, X. B., Yan, P., & Zhu, H. T. (2009). Telomere length shortens with age in Chinese alligators (*Alligator sinensis*). *Journal of Applied Animal Research*, *36*(1), 109-112.

Zhang, Q., Han, X., Hao, X., Ma, L., Li, S., Wang, Y., & Du, W. (2018). A simulated heat wave shortens the telomere length and lifespan of a desert lizard. *Journal of Thermal Biology*, *72*, 94-100.
